# Cross-linguistic and acoustic-driven effects on multiscale neural synchrony to stress rhythms

**DOI:** 10.1101/2023.12.04.570012

**Authors:** Deling He, Eugene H. Buder, Gavin M. Bidelman

## Abstract

We investigated how neural oscillations code the hierarchical nature of stress rhythms in speech and how stress processing varies with language experience. By measuring phase synchrony of multilevel EEG-acoustic tracking and intra-brain cross-frequency coupling, we show the encoding of stress involves different neural signatures (delta rhythms = stress foot rate; theta rhythms = syllable rate), is stronger for amplitude vs. duration stress cues, and induces nested delta-theta coherence mirroring the stress-syllable hierarchy in speech. Only native English, but not Mandarin, speakers exhibited enhanced neural entrainment at central stress (2 Hz) and syllable (4 Hz) rates intrinsic to natural English. English individuals with superior cortical-stress tracking capabilities also displayed stronger neural hierarchical coherence, highlighting a nuanced interplay between internal nesting of brain rhythms and external entrainment rooted in language-specific speech rhythms. Our cross-language findings reveal brain-speech synchronization is not purely a “bottom-up” but benefits from “top-down” processing from listeners’ language-specific experience.

**Highlights:**

- Neural oscillations at delta and theta bands synchronize with stress and syllable rhythms.
- Hierarchical delta-theta phase coupling mirrors speech rhythm hierarchy.
- Language experience shapes multiscale brain-to-speech entrainment.
- Optimized brain-to-speech synchronization occurs at natural stress (2 Hz) and syllable (4 Hz) rates.
- Amplitude cues dominate the neural oscillatory encoding of stress rhythm.

## 1. INTRODUCTION

A growing number of brain imaging studies suggest that speech is processed at multiple temporal windows operated by a set of neuronal oscillators whose frequencies are tuned to relevant features of the acoustic-linguistic signal (Ding et al., 2016; Ghitza, 2011; Gross et al., 2013; Hyafil et al., 2015; Kösem & Van Wassenhove, 2017; Poeppel, 2003; Rimmele et al., 2023; Teng et al., 2017). The oscillations associated with speech are spectrally distributed in the gamma (> 30 Hz), theta (4 - 8 Hz), and delta (1- 3 Hz) frequency bands of the EEG, roughly corresponding with the time spans of phonemic, syllabic, and supra-syllabic units. Presumably, the processing of speech might be realized through the phase alignment of brain oscillations to the speech amplitude envelopes, which segment/parse the continuum speech signals into linguistic representations (Doelling et al., 2014; Ghitza, 2012; Luo & Poeppel, 2007).

Such brain-to-speech synchronization is especially significant in terms of coding syllable rhythm. Theoretical and empirical work suggests brain activity imposes a constraint on processing such that auditory perception is optimized when theta band (4-8 Hz) oscillations coincide with the range of natural syllable rates (Ghitza, 2012; Houtgast & Steeneken, 1985; Luo & Poeppel, 2007; Poeppel & Assaneo, 2020). For instance, speech intelligibility is severely degraded with low-pass filtering below 2 Hz and is only marginally improved by adding modulations above 8 Hz (Drullman et al., 1994). Moreover, cortical-acoustic entrainment and intracranial auditory-motor coherence is enhanced at frequencies close to the dominant syllable rhythm which has been empirically found to be 4-5 Hz across languages (Assaneo & Poeppel, 2018; He et al., 2023). However, whether there are also optimal supra-syllabic frequencies within lower-frequency delta neural oscillations has not been explicitly tested, though several studies have begun to examine delta-neural entrainment.

Cycles of delta oscillations often align with repetitive complex sounds including frequency-modulated complex tones (Henry & Obleser, 2012), digit strings (Rimmele et al., 2021), noise-vocoded speech (Bröhl & Kayser, 2021), prosodic or lexical phrases (Cogan & Poeppel, 2011; Gross et al., 2013; Keitel et al., 2018; Lo et al., 2022), and sentences (Lu et al., 2022). However, the particular sound elements that entrain delta oscillation remain elusive, being variably attributed to “intonation, prosody, and phrases” (Boucher et al., 2019; Ghitza, 2011; Giraud & Poeppel, 2012; Gross et al., 2013; Rimmele et al., 2021). Moreover, prominent theories in language processing, such as the asymmetric sampling in time (AST) or TEMPO hypotheses, have overlooked the potential hierarchical role of delta oscillations (Ghitza, 2011; Ghitza & Greenberg, 2009; Hickok & Poeppel, 2007; Poeppel, 2003). This has led to a growing debate, with some arguing ongoing delta oscillations modulate theta activity (Gross et al., 2013; Lakatos et al., 2005), and others asserting the master role of the theta oscillator (Ghitza, 2011, 2013). These discrepancies pinpoint the necessity for a more integrated exploration of delta oscillations, which have generally been overlooked in the literature.

However, there are several prominent suprasegmental features of speech that might be optimally coded by delta brain oscillations. One important feature that creates a natural hierarchy in speech is stress. Specifically, stress foot^1^ is a supra-syllabic unit that organizes a group of syllables by assigning emphasis on the stressed syllable (Hogg et al., 1987; Leong, 2012; Selkirk, 1980). Approximately 85% of English words begin with the first syllable stressed in English (Cutler & Carter, 1987), which is mostly signaled by higher amplitude and longer duration (Fry, 1955; Greenberg et al., 2003), and to a lesser extent, pitch (Arvaniti, 2009; Greenberg, 1999; Kochanski et al., 2005; Silipo & Greenberg, 1999, 2000). Importantly, acoustic research illustrates that English simultaneously carries syllable and stress foot rhythms in the speech signal, represented by frequency-specific amplitude envelopes that closely correspond to theta and delta brain oscillations (Greenberg et al., 2003; Leong, 2012; Leong et al., 2014; Tilsen & Arvaniti, 2013).

The hierarchical nature of stress assignment also creates the opportunity for nesting of different speech elements. For example, while the dominant syllable rhythm is at 4 - 5 Hz across languages, the rhythm of stress foot in English is centered at half this speed, nominally around 2 Hz (Ding et al., 2017; Greenberg, 1999; Greenberg et al., 1996; Tilsen & Arvaniti, 2013; Tilsen & Johnson, 2008). Indeed, faster syllable rhythms are embedded into slower stress foot constituents, creating hierarchical nesting (Goswami & Leong, 2013; Leong, 2012). Such hierarchy is quantitatively illustrated by cross-frequency phase coupling seen in different acoustic constituents of the speech amplitude envelope. For example, Goswami and Leong (2013) showed a phase hierarchy relationship, where the ridge in the stress foot envelope always aligns with the stressed syllable and away from the unstressed syllable. Such hierarchy constrains stressed syllables to occur only in certain phases of the stress foot envelope (Leong, 2012). This is of particular interest given that the relationship between delta and theta brain oscillations may provide one such mechanism that mirrors this hierarchical structure of speech. However, there remains an empirical gap on how multiscale brain oscillations lock to the hierarchical properties of stress rhythms. To our knowledge, whether such stress foot-syllable hierarchy seen in the speech signal is reflected neurobiologically in delta-theta brain rhythm coupling remains to be tested.

The hierarchical nature of stress also affords the opportunity to explore cross-language differences in delta-theta mechanisms of speech processing. Yet, the specifics of brain oscillatory dynamics that vary among speakers with distinct language backgrounds remains largely unexplored. Still, some studies show cortical oscillations reliably track the syllabic amplitude envelope even in a foreign or unintelligible language (Ding et al., 2016; Zou et al., 2019). However, this brain-speech tracking appears to falter at the suprasyllabic level, and foreign listeners struggle to understand the linguistic content conveyed by delta band activity (Blanco-Elorrieta et al., 2020; Ding et al., 2016). These findings suggest the possibility of language-specific tuning of brain oscillatory dynamics, particularly pertaining to processing at the suprasyllabic level.

Indeed, in the context of stress encoding, it is reasonable to assume that cross-linguistic differences in delta band oscillations might occur in native English vs. nonnative speakers owing to the relative importance of stress in English vs. other languages. In particular, a comparison between English and Mandarin Chinese listeners could elucidate experience-dependent changes in stress-related brain processing given the distinctive prosodic features in each language (Hogg et al., 1987; Jongman et al., 2006). Supporting this cross-language design, behavioral and EEG studies have in fact shown that intensity is a less reliable cue for Mandarin listener’s perception of English stress given the lesser importance of this cue in their native language (Mandarin) (Chrabaszcz et al., 2014; Chung & Bidelman, 2016). Thus, one primary objective herein was to further characterize such cross-language differences in oscillatory stress processing.

The current study aimed to examine delta (syllable level) and theta (stress foot level) oscillations in terms of how these neural correlates of rhythmic stress processing vary with language experience and acoustic modulations. We analyzed multilevel EEG-acoustic phase synchrony and intra-brain cross-frequency coupling while English and Chinese listeners perceiving various rhythmic stress patterns. We hypothesized that brain oscillations in the delta and theta bands would concurrently synchronize to stress and syllable rhythms, given their putative role in coding these properties of speech. Furthermore, we hypothesized this brain-stress synchronization might be enhanced by the dominant natural stress patterns (e.g., amplitude-signaled high salient stress rhythm at 2 Hz) in English speech. We further posited the acoustic phase hierarchy between stress foot and syllable rhythms (Leong, 2012; Leong et al., 2014) would be paralleled in the brain (EEG) as enhanced coupling of delta-theta oscillations. Lastly, we hypothesized Chinese speakers might have reduced neural responses coding stress rhythm (given the relative unimportance of stress in their native Mandarin), yet maintain neural entrainment to syllable rhythm—which is likely more discernable to even non-native speakers.

## 2. MATERIALS & METHODS

### 2.1 Participants

The study included N = 34 young adults recruited from the University of Memphis student body and Greater Memphis area. N = 17 were native speakers of American English (7 males and 10 females) and N = 17 were native speakers of Mandarin Chinese (7 males and 10 females). The two groups were closely matched in age (English: 24.9 ±4.6 years; Chinese: 27.3 ±3.5 years), years of education (English: 18.5 ±3.76 years; Chinese: 20.8 ±2.56 years), and musical training (English: 6.9 ±6.4 years; Chinese: 5.3 ±8.0 years). The majority of participants were right-handed (English: 60% ± 60%; Chinese: 61% ±44%), as evaluated using the Edinburgh Handedness Inventory (Oldfield, 1971). All participants had normal hearing sensitivity, defined as pure tone thresholds of ≤ 25 dB HL at octave frequencies from 500 Hz to 8000 Hz in both ears. There was no history of speech, language, or neuropsychiatric disorders reported among participants.

We used a language history questionnaire to assess language background (e.g., Li et al., 2006). Our inclusion criteria for native Mandarin speakers were consistent with prior cross-language EEG studies (Bidelman et al., 2011; Blanco-Elorrieta et al., 2020; Chung & Bidelman, 2016). Chinese listeners were born and raised in China, with first exposure to English beginning in school around the age of 7.8 ±2.63 years. They resided in the United States (US) during the experiment, with a duration of stay of 4.9 ±3.61 years. Their self-reported English proficiency was moderate to high (4.9 ±1.09, with a score of 7 indicating native-like proficiency). Each reported using native Mandarin approximately 59% ±17% of their daily communication. Two Mandarin speakers who also speak Cantonese were excluded from the study because of potential confounds related to Cantonese listeners’ advantages in stress perception (Choi, 2021). All participants provided written informed consent in accordance with a protocol approved by the University of Memphis Institutional Review Board and received compensation for their involvement.

### 2.2 EEG Stimuli

Audio tokens of the syllables ‘ba’ and ‘ma’ were recorded by a male talker (2^nd^ author) spoken in isolation with natural and similar loudness and pitch. These syllables were similar to stimuli used in our previous study on neural speech entrainment (He et al., 2023). Each syllable underwent temporal compression to a fixed duration of 120 ms and then concatenated to form pairs (e.g., ‘baba’) in Praat (Boersma & Weenink, 2013). We then separately manipulated syllable amplitude and duration to create four different stress tokens (e.g., ‘BAba’) where the first syllable was stressed, conforming to the trochaic foot (**Figure 1**).

**Figure 1:**
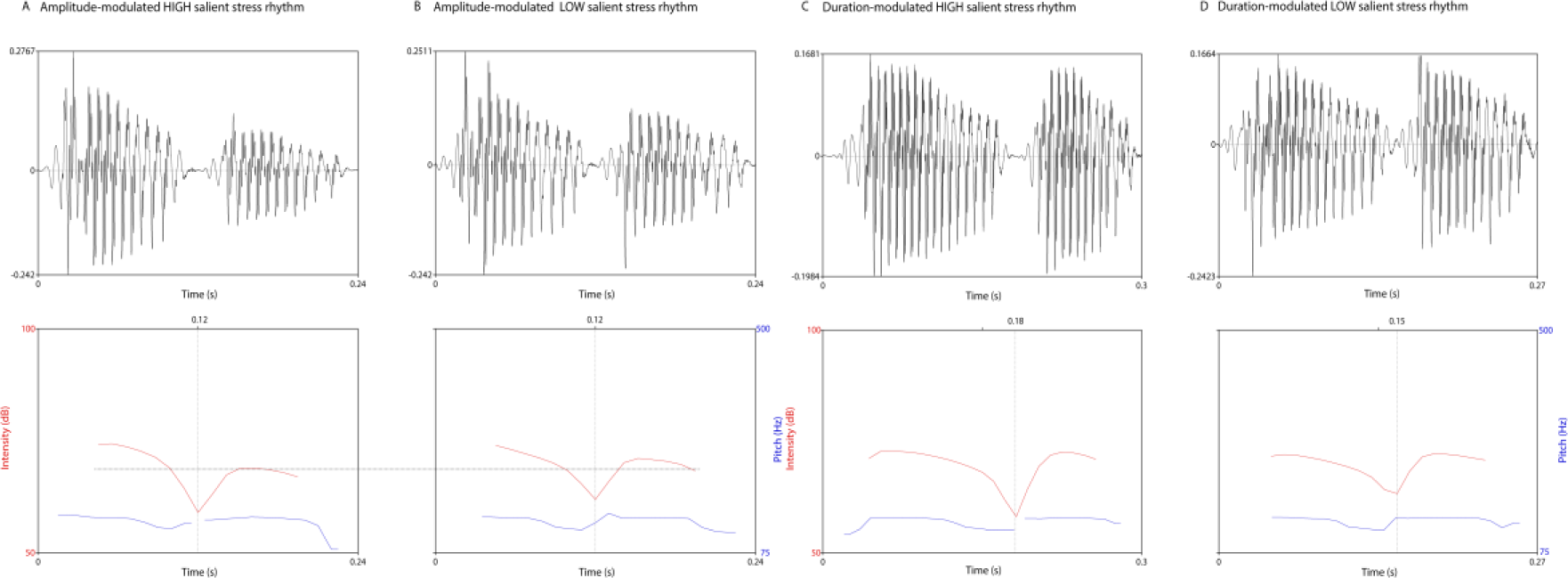
Examples of stress stimuli used in the EEG study (i.e., a trochaic stress token). Disyllables were modulated by amplitude envelope (A & B) and syllable duration (C & D) with a high or low stress salience, respectively. black = speech sound stimulus; red = intensity contour; blue = pitch contour; dash lines mark the comparison between stressed and unstressed syllables in amplitude or duration.

#### Amplitude-signaled stress pattern

(Fig. 1 A & B). High salient tokens were characterized by a 50% higher amplitude between stressed and unstressed syllables, while low stress tokens were reduced to a 25% contrast. However, each syllable maintained a uniform 120 ms duration across both salience levels.

#### Duration-signaled stress pattern

(Fig. 1 C & D). Similar with the amplitude condition, the duration contrast between stressed and unstressed syllables was marked at 50% and 25% for high and low saliences, respectively. High salient tokens featured 180 ms stressed syllables paired with 120 ms unstressed syllables, while low salient tokens contained 150 ms stressed syllables alongside 120 ms unstressed counterparts, all maintaining uniform amplitude.

Finally, each stress disyllable was concatenated (by inserting silence) to generate a continuous speech train of 6 seconds at different rates of 1, 2, and 3 Hz. These rates were chosen because English stress rhythm typically unfold with a nominal rate around 2 Hz (Dauer, 1983; Leong, 2012; Tilsen & Arvaniti, 2013). Altogether, we generated 12 stimulus conditions, each featuring a distinct stress pattern due to manipulation of acoustic cue (amplitude or duration), salience (high or low), and rhythm (1, 2, and 3 Hz).

### 2.3 Data acquisition and preprocessing

During electrophysiological recordings, participants comfortably reclined in front of a PC monitor and performed speech perception tasks in an electro-acoustically shielded booth (Industrial Acoustics Company). Binaural auditory stimuli were presented at 84 dB SPL through ER-2 insert earphones (Etymotic Research). Stimulus intensity was calibrated using a Larson-Davis SPL meter measured in a 2-cc coupler (IEC 60126). The presentation of stimulus and task instructions was managed by MATLAB 2013 (The MathWorks, 2013) directed to a TDT RP2 signal processing interface (Tucker-Davis Technologies). Participants were instructed to listen to the speech streams and identify whether they heard “STRONG weak” or “weak STRONG” syllable sequences using the keyboard (labeled as ‘AAbb’ or ‘aaBB’). There were no time limits for behavioral responses. The subsequent trial commenced after the listener’s response. Each stress stimulus condition comprised 10 trials (each 6 s). The presentation order of the conditions was randomized both within and across participants.

Continuous EEG signals were recorded using Ag/AgCl disc electrodes placed at the mid-hairline and referenced to linked mastoids (A1/A2), with the mid-forehead serving as the ground. This single-channel montage is highly effective in recording entrained, auditory neural responses to speech (He et al., 2023) generated from auditory cortex (Bidelman et al., 2013; Picton et al., 1999). Inter-electrode impedance was maintained < 10 kΩ. Continuous EEGs were digitized at a sampling rate of 1000 Hz using SynAmps RT amplifiers (Compumedics Neuroscan) and an online passband filter of 0- 400 Hz.

Subsequent preprocessing was conducted using a customized MATLAB script. To focus on the slow electrophysiological activities, neural signals were further passband filtered (0.9-30 Hz; 10^th^ order Butterworth). First, EEGs were segmented into individual 6-s epochs^2^–conforming to the length of the audio stimulus—and concatenated, resulting in 60 s of EEG data per condition. To minimize eye blink artifacts, we applied a wavelet-based denoising algorithm to the continuous EEGs (Khatun et al., 2016). **Figure 2A** shows examples of one trial of EEG data corresponding to delta (stress) and theta (syllable) neural responses from the English group for the 1, 2, and 3 Hz stress rates, respectively. **Figure 2 B** presents the corresponding spectrum of 60-s of continuous EEG data.

**Figure 2:**
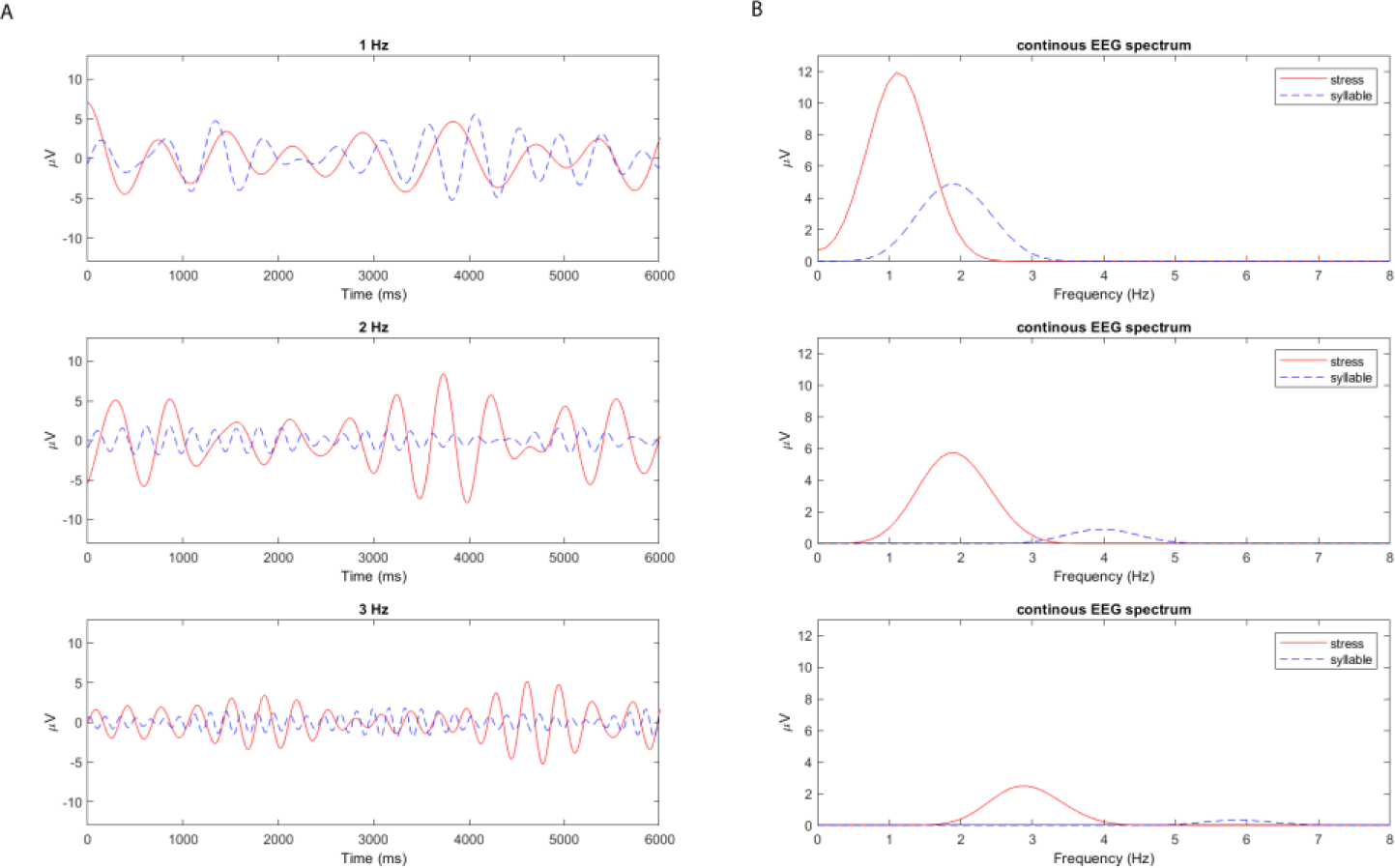
Continuous EEGs show phase-locking to stress and syllable rhythms. Shown here is an example from EEGs of the initial trial (6 s) that were averaged across English individuals for the amplitude high stress condition. (A) neural phase coupling dynamic represents the phase hierarchy in speech rhythms. (B) Spectrum analysis quantifies the intensity of each frequency of interest. Red = frequencies of stress rhythm; blue = frequencies of syllable rhythm.

### 2.4 Electrophysiological data analysis

Phase Locking Value (PLV) and n:m Phase Synchronization Index (nmPSI) are bivariate time-series measures that quantify the degree of phase synchronization between two oscillators or time series. PLV computes the phase synchrony of two time series (e.g., acoustic and EEG signals) at a singular frequency (Assaneo & Poeppel, 2018; He et al., 2023; Lachaux et al., 1999). In contrast, nmPSI evaluates the cross-frequency phase coupling between two oscillators with distinct frequencies described by n and m (e.g., delta and theta frequency bands of EEG signals), where n:m is an integer relation (Leong et al., 2017; Rosenblum et al., 1998; Schack & Weiss, 2005). Conceptually, both PLV and nmPSI capture the temporal consistency in phase difference (and, conversely, the coherence) between two signals. Their resulting values range from 0 (no synchronization) to 1 (complete synchronization). PLV and nmPSI were computed using the following formulas:

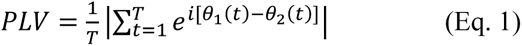

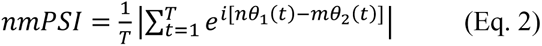

Here, t denotes the discretized time, T is the total number of time points, and θ1(t) and θ2(t) are the Hilbert phases of the first and second signals, respectively.

The current study assessed synchronization between neural and acoustic speech signals using PLV at frequencies corresponding to stress rhythm (i.e., 1, 2, and 3 Hz) and syllable rhythm (i.e., 2, 4, and 6 Hz), respectively. This results in PLV_Stress_ representing brain-acoustic synchronization at the stress level and PLV_syllable_ reflecting brain-acoustic synchronization at the syllable level. We measured nmPSI to quantify the cross-frequency coupling within the brain’s theta and delta frequency bands, corresponding to the alignment of nested syllable and stress rhythms unfolding at a 2:1 ratio. Specifically, frequency-specific neural signals and acoustic inputs were computed by applying passband filters around the frequencies of interest (±0.5 Hz) (see Fig. 2). The phase was extracted as the imaginary part of the signal’s Hilbert transform. PLV was then computed between the EEG signal and acoustic stimulus waveform within each narrow frequency band and averaged over time per individual according to Equation 1. In contrast, nmPSI was computed by bandpass filtering the EEG data into two separate bands (i.e., m = 1, 2, 3 ±0.5 Hz; n = 2, 4, 6 ±0.5 Hz) to isolate phase-locked responses to the stress (m) and syllable (n) rhythm in the brain at a 2:1 ratio. To reduce noise in the metric, we quantified nmPSI in a moving window (6 sec; overlap ratio of 0.3) and averaged across windows for each condition according to Equation 2. To establish the noise floor of our PSI metric, we applied this identical analysis to our previous EEG data evoked by similar syllable trains but devoid of any stress patterns (e.g., ‘ba-ba-ba…’) (He et al., 2023).

### 2.5 Statistical analysis

We conducted four-way mixed model analyses of variance (ANOVAs) in R (version 1.3.1073; ‘lme4’ package; Bates et al., 2015) to assess whether multi-scale brain-to-speech synchrony and cross-frequency coupling within the brain differed due to the acoustic stress patterns and language experience by measuring PLV_Stress_, PLV_syllable_, and nmPSI. The model included within-subject factors of the stress cue (2 levels; amplitude vs. duration), stress salience (2 levels; high vs. low), and stress rate (3 levels; 1, 2, and 3 Hz) and a between-subject factor of group (2 levels; English vs. Chinese); subjects served as a random factor [e.g., PLV∼ cue*salience* rate*group+(1|sub)]. We used Tukey post hoc tests to correct for multiple comparisons. Given our *a priori* hypothesis regarding potential enhancements of synchronization at 2 Hz (nominal English stress rhythm), following the initial omnibus ANOVA, we examined contrasts for nmPSI and PLV_Stress_ between 2 Hz versus the other stress rates. Similar contrast was conducted for PLV_syllable_ between the nominal rate of 4 Hz versus others.

Furthermore, to assess associations between neural-neural and neural-acoustic synchrony measures, we used repeated measures correlations (rmCorr; Bakdash & Marusich, 2017). Unlike conventional correlations, rmCorr accounts for non-independence among each listener’s observations and measures within-subject correlations by evaluating the common intra-individual association between two measures. Preliminary diagnostics (quantile–quantile plot and residual plots) were used to validate normality and homogeneity assumptions. Behavioral data from the EEG task (i.e., percentage of correctly perceived stress patterns) were rationalized arcsine transformed (Studebaker, 1985). A priori significance level was α = 0.05. Effect sizes are presented as *n*^2^.

## 3. RESULTS

Our behavioral task was primarily designed to keep subjects attentive and awake rather than assess stress perception, per se. Indeed, correct percent performance showed no group differences and results approached chance level (see Supplemental material; Fig. S1).

### 3.1 Brain to speech tracking at the *stress level*

**(PLV_Stress_)** We examined how neural oscillations phase lock to the external (i.e., acoustic) stress rhythms at rates of 1, 2, and 3 Hz (**Fig. 3**). An ANOVA conducted on PLV_Stress_ revealed significant main effects for group (*F_1,32_* = 4.69, *p* = 0.038, 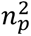 = 0.13), stress rate (*F_2,352_* = 10.03, *p* = 0.0001, 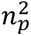 = 0.05), cue (*F_1,352_* = 4.02, *p* = 0.046, 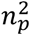 = 0.01), along with a two-way stress cue * salience interaction (*F_1,352_* = 8.65, *p* = 0.003, *n*^2^ = 0.02). Notably, English listeners demonstrated stronger brain-to-acoustic stress tracking than native Chinese speakers. The rate effect was attributed to stress rhythms at 1 and 2 Hz eliciting greater PLV_Stress_ (*p_1 vs. 3 Hz_* = 0.001; *p2 vs. 3 Hz* = 0.0001) compared to 3 Hz. The interaction of cue*salience arose from enhanced PLV_Stress_ for amplitude-(*p* < 0.001) compared to duration-signaled high salient stress, suggesting a neural preference of amplitude cues for both groups. Also, when stress was signaled by duration, we found higher PLV_Stress_ for low compared to high stress salience (*p* < 0.001). These results highlight the differences in exogenous neural-acoustic synchronization across individuals’ language experience and stress rhythm rates.

**Figure 3:**
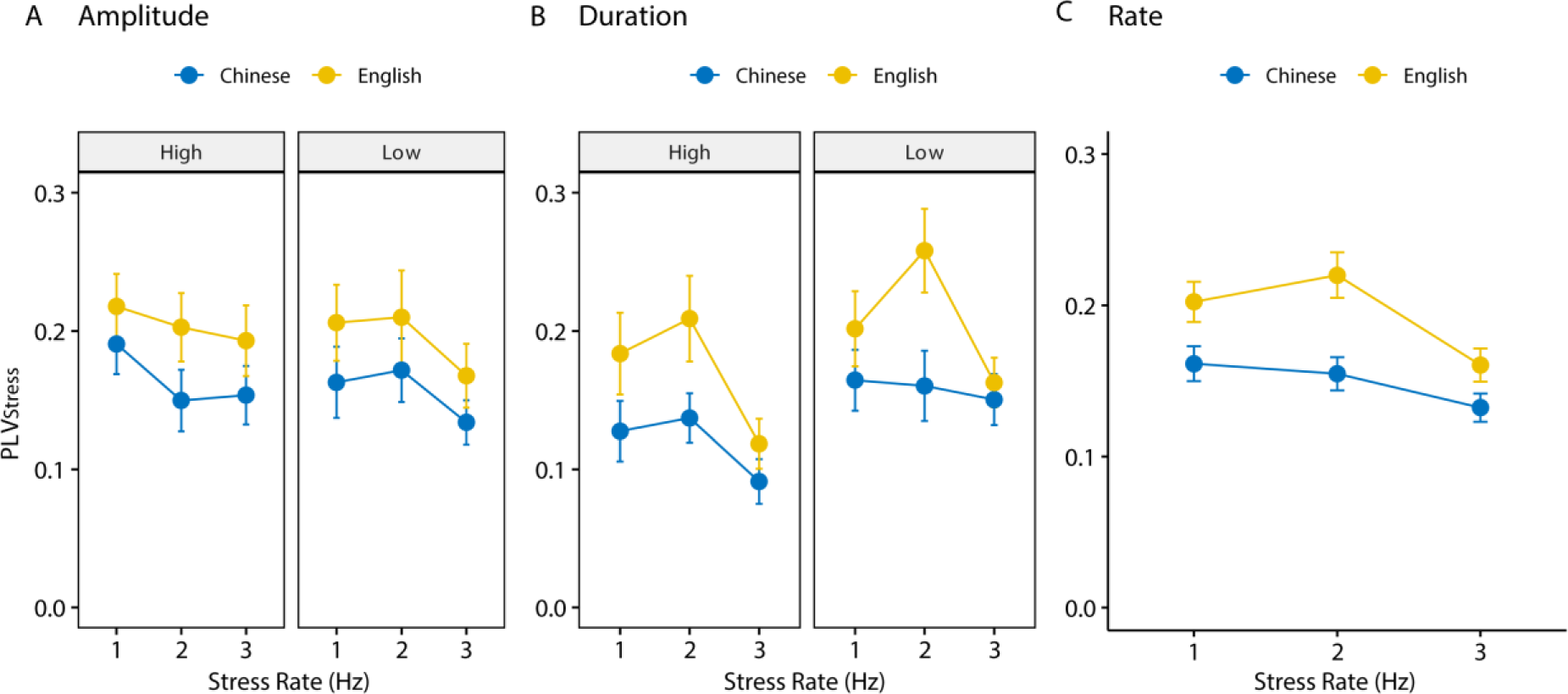
Brain oscillations synchronize to the rate of *stress rhythms*. Cross-linguistic PLV_Stress_ comparisons by stress rate and salience signaled by (A) amplitude envelope and (B) syllable duration. (C) English differed from Chinese speakers across rates and exhibited PLV_Stress_ enhancement at 2 Hz**—**corresponding to the natural stress rate found in English. Panel C outlines the main effects of rate, arrogating data across cue and salience conditions. PLV_Stress_ refers to neural-to-stress rhythm phase locking; errorbars = ±1 s.e.m.

Motivated by the predominant rate of 2 Hz in natural English stress rhythms (Dauer, 1983; Leong, 2012; Tilsen & Arvaniti, 2013) and the inverted-V rate pattern depicted in Figure 3, we conducted an *a priori* contrast of 2 Hz against other rates by group and stress cue. Our assumption was confirmed in that PLV_Stress_ peaked at 2 Hz exclusively for English speakers (*p* = 0.0001) under duration-modulated stress. Interestingly, this enhancement was absent for Chinese whose native language does not include English-based stress patterns (*p_amplitude_* = 0.98; *p_duration_* = 0.373). Our results demonstrate an enhancement of speech-to-brain phase-locking (PLV_Stress_) at the frequency inherent to natural English stress rhythm (2 Hz), that is also shaped by individuals’ language exposure.

### 3.2 Brain to speech tracking at *syllable level*

**(PLV_syllable_)** As our stimuli simultaneously carried stress and syllable rhythms organized as hierarchical tiers, we next proceeded to test the extent to which neural oscillations phase lock to the acoustic syllable rates^3^ of 2, 4, and 6 Hz, which are 2-times faster than stress rhythms. An ANOVA conducted on PLV_syllable_ revealed a main effect of stress rate (*F_2,352_* = 5.93, *p* = 0.003, *n*^2^ = 0.03) and two-way interactions of stress cue * group (*F_1,352_* = 5.79, *p* = 0.017, *n*^2^ = 0.02) and stress cue * rate (*F_2,352_* = 5.53, *p* = 0.004, *n*^2^ = 0.03) (**Fig. 4**). Pairwise comparisons revealed English speakers had higher PLV_syllable_ than Chinese speakers under amplitude-(*p* = 0.05) but not duration-signaled stress. This group difference was further Tukey pairwise compared by rates and was only significant at a syllable rate of 4 Hz under amplitude-indicated stress patterns (*p* = 0.005) motivated by the cue* rate interaction. For English speakers, amplitude-signaled stress rhythm had a stronger PLV_syllable_ than duration-signaled stress (*p* = 0.029), consistent with PLV_Stress_. The stress cue * rate interaction was driven by stronger PLV_syllable_ for duration cues at 3 Hz compared to other rates (*p_3 vs. 2 Hz_* < 0.001; *p_3 vs. 2 Hz_* = 0.001). Additionally, at 3 Hz, duration cues evoked higher PLV_syllable_ than amplitude cues.

**Figure 4:**
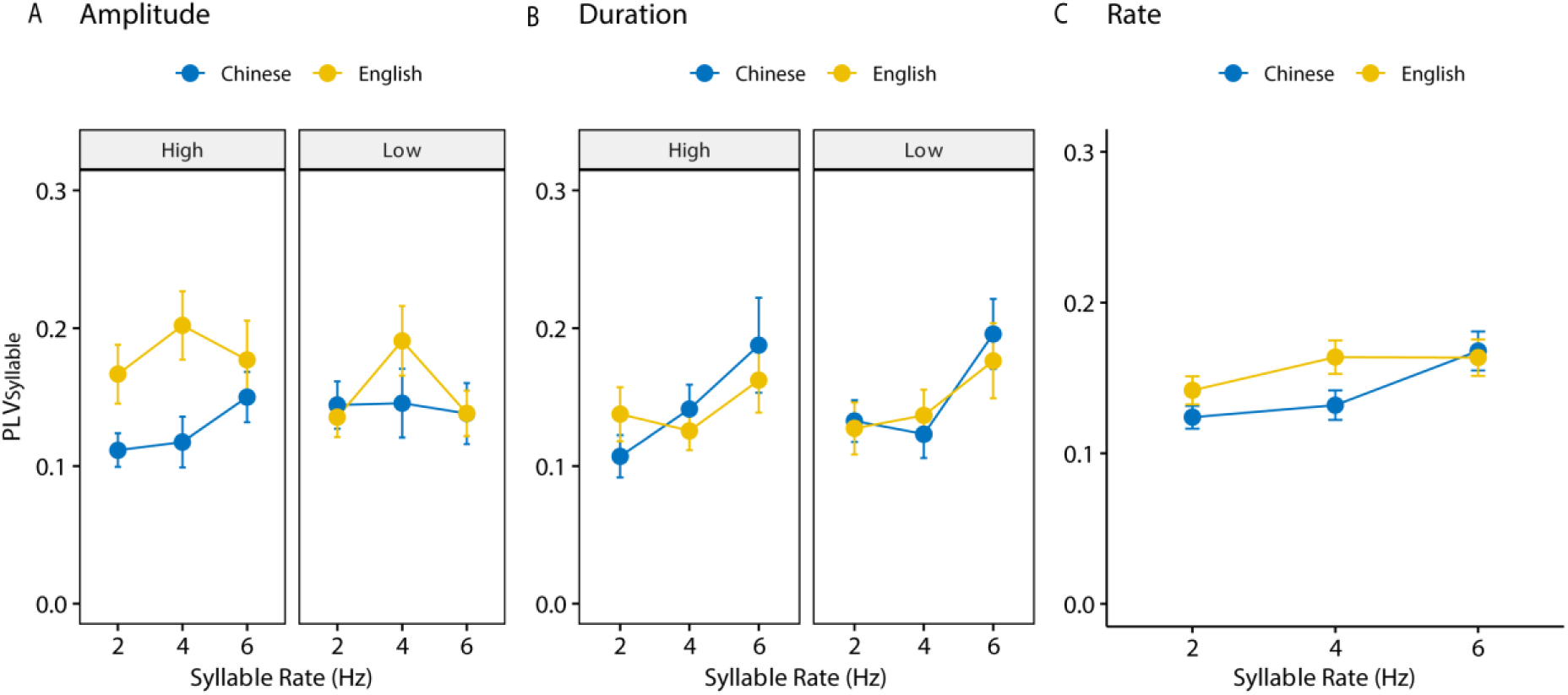
Brain oscillations phase lock to the rate of *syllable rhythms.* Cross-linguistic PLV_syllable_ comparisons by stress rate and salience signaled by (A) amplitude envelope and (B) syllable duration. (C) PLV_syllable_ enhancement at syllable rhythm of 4 Hz, matching the center syllable rate across many languages, was exclusively observed in English speakers, not Chinese. Panel C outlines the main effects of rate, arrogating data across cue and salience conditions. PLV_syllable_ refers to neural-to-syllable rhythm phase locking; error bars = ±1 s.e.m.

Consistent with PLV*Stress*, we performed an *a priori* contrast involving the 4 Hz-syllable rate (that is, 2 Hz-stress rate) versus other rates (i.e., 2 and 6 Hz syllable rates) by group and stress cue. We observed an enhanced PLV_syllable_ at 4 Hz (*p* = 0.011) only in English speakers for amplitude cues. Notably, 4 Hz closely aligns with the mean syllable rate in English (Goswami & Leong, 2013; Greenberg et al., 2003; Tilsen & Johnson, 2008) and many other languages, including Chinese (Ding et al., 2017). Surprisingly, this preferred syllable rate of 4 Hz in natural speech failed to result in neural coding enhancement for Chinese speakers, possibly due to their limited brain tracking at the stress level (cf. Fig. 3). Together, these findings validate that neural-acoustic synchronization is modulated by syllable rates and is enhanced when aligned to the natural syllable rate. They also confirm the presence of *stress-level* brain tracking.

### 3.3 Cross-frequency coupling within the brain

**(nmPSI)** Figure 5 illustrates delta-theta phase coupling *within the brain* as measured by nmPSI. Results yielded significant main effects, including group (*F_1,32_* = 90.42, *p* < 0.0001, 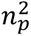 = 0.74), stress rate (*F_2,352_* = 157.82, *p* < 0.0001, *n*^2^ = 0.47), cue (*F_1,352_* = 91.03, *p* < 0.0001, 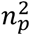 = 0.21), and salience (*F_2,352_* = 87.88, *p* < 0.001, *n*^2^ = 0.20). Post hoc analysis indicated peak nmPSI occurred at a stress rate of 1 Hz, declining gradually with faster rates (*all p* < 0.0001) for both groups. Moreover, English speakers had greater nmPSI than Chinese speakers across all stress rates (Fig. 5C). In addition, we found a significant three-way interaction of cue * salience * group (*F_1,352_* = 75.90, *p* < 0.0001, 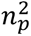 = 0.18), which was attributed to English speakers having stronger nmPSI relative to Chinese speakers for low salient stress stimuli (*p_amplitude_* < 0.0001; *p_duration_* = 0.0026) (**Figs. 5A and B**, right panels). However, no group differences were observed for more salient (i.e., high) stress stimuli—true for both stress cues (**Figs. 5A & B**, left panels). Generally speaking, both Chinese and English speakers exhibited nmPSI above baseline, indicating internal delta-theta coupling represented the stress-syllable hierarchy of our stimuli above what would be expected by random variation alone. Crucially, this phase hierarchy in the brain was enhanced in English listeners who possess extensive experience in a language specifically structured by stress patterns.

**Figure 5:**
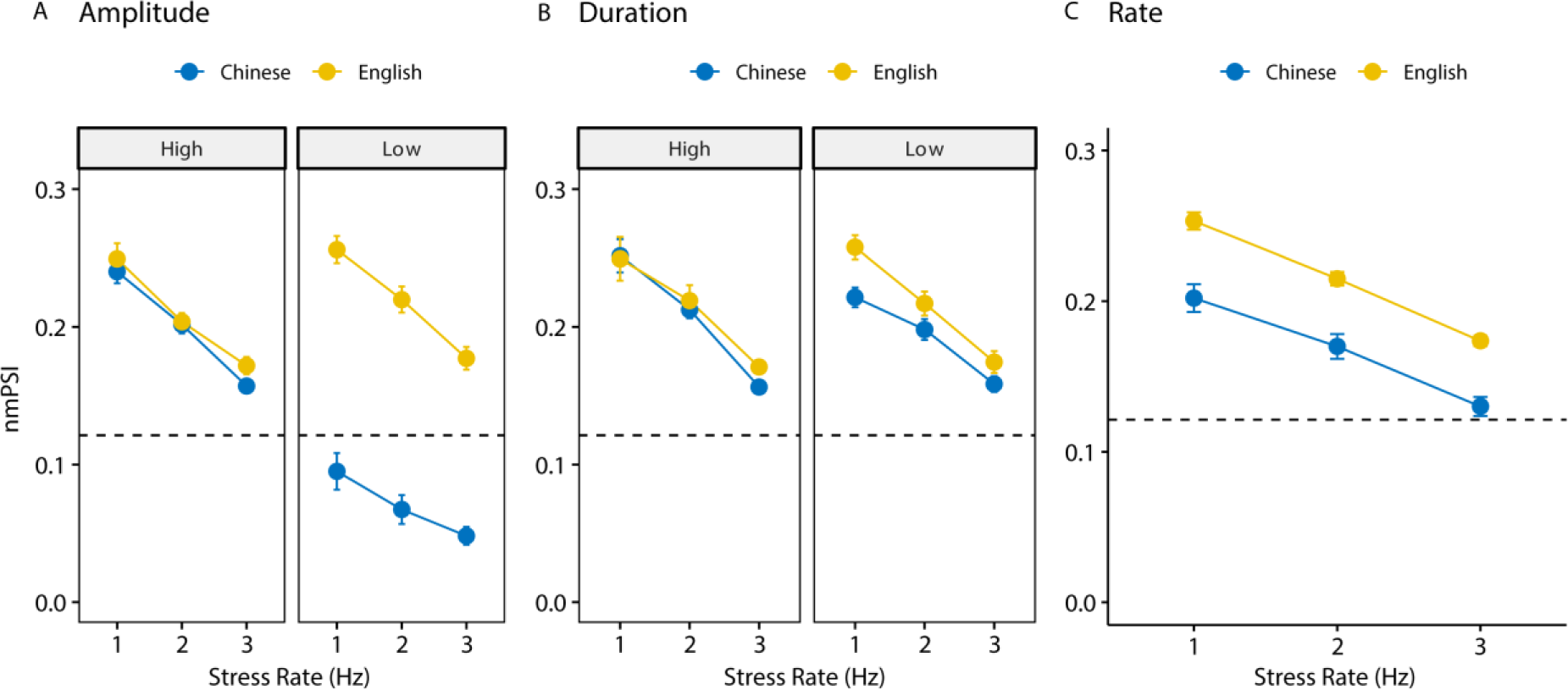
Phase coupling of delta-theta neural oscillations represents the phase hierarchy of stress-syllable rhythms. Cross-language comparisons of nmPSI as a function of stress rate and salience modulated by (A) amplitude envelope and (B) syllable duration (C) Significant group differences in nmPSI across stress rhythm rate with a dataset arrogated across cue and salience conditions. Dashed lines = nmPSI baseline computed for stress-free syllable trains from He et al. (2023). error bars = ±1 s.e.m.

Figure 6 illustrates the data broken down by stress cue (amplitude vs. duration), cue salience, and group to emphasize language-specific cue differences. Pairwise comparisons revealed that nmPSI differences in stress cue and salience were only observable for the Chinese group (Fig. 6B), where high salient stress resulted in higher nmPSI compared to low salience for both cues (*p_amplitude_* < 0.0001; *p_duration_* = 0.043). Also, under low stress salience, duration-related stress had higher nmPSI than amplitude stress (*p* < 0.0001) within Chinese. Critically, there were no significant nmPSI differences due to acoustic stress cue nor salience for native English speakers (Fig. 6A; 3-way ANOVA: *pcue* = 0.727; *psalience* = 0.196). These findings suggest that English listeners coded and constructed consistent neural coherence that was equally robust across varying acoustics. That is, English speakers’ neural responses effectively entrained to the hierarchical stress rhythms even in scenarios where the aural information cuing stress was relatively weak in perceptual salience. In stark contrast, Chinese listeners’ hierarchical entrainment nmPSI was more susceptible to acoustic variations signaling stress patterns.

**Figure 6:**
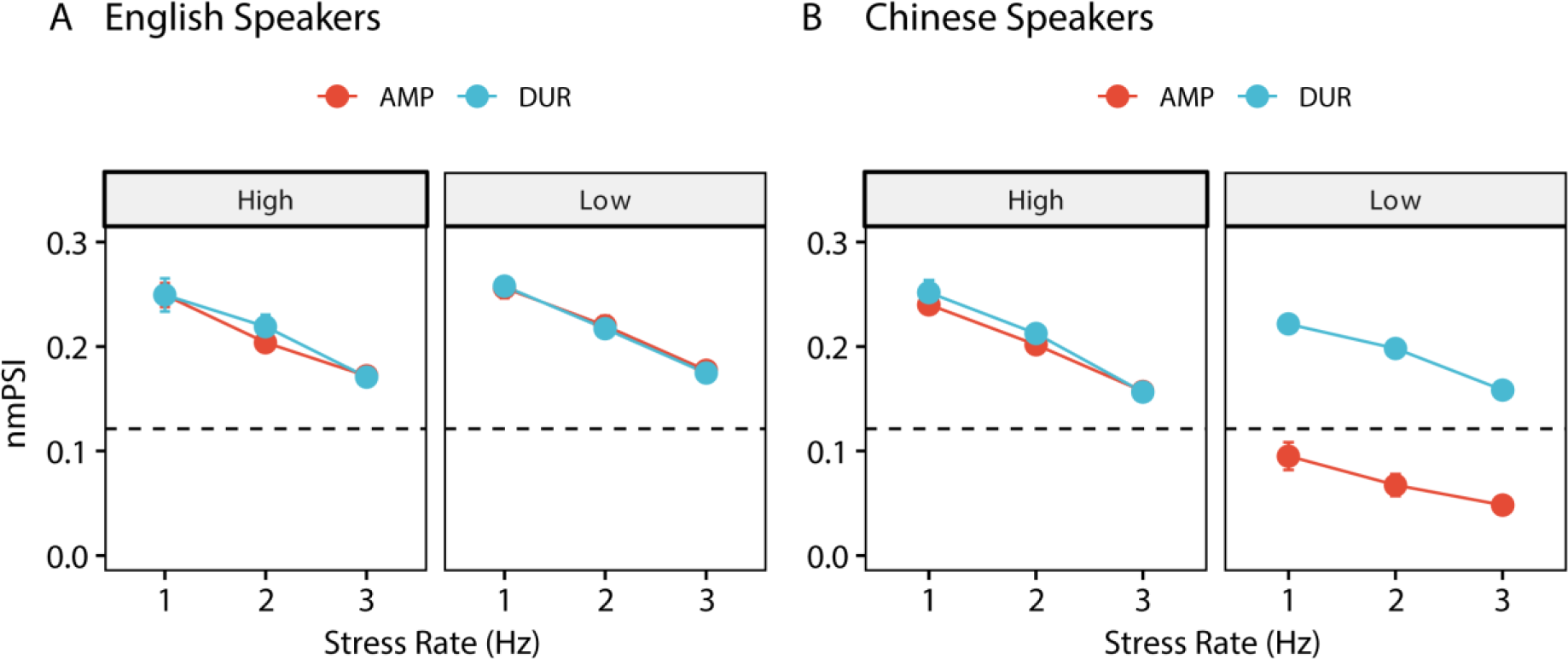
Effects of acoustic stress cue and salience on cross-frequency coupling. (A) nmPSI in English was invariant to acoustic stress manipulations (B) Contrastively, Chinese listeners’ nmPSI was more prone towards high salience and duration cue, suggesting stronger coupling of their nested brain oscillations in these conditions. Dashed lines = nmPSI baseline computed for stress-free syllable train perception. error bars = ±1 s.e.m.

### 3.4 Correlations

To explore the association between internal brain cross-frequency synchronization and external brain-speech synchronization, we conducted within-subject correlations using rmCorr for all the feasible pairwise variables (i.e., nmPSI, PLV_Stress_, PLV_syllable_). Figure 7 depicts a positive correlation between nmPSI and PLV_Stress_ for English (*r* = 0.23, *p* = 0.002) but not Chinese (*r* = 0.07, *p* =0.352) speakers. English individuals exhibiting stronger internal cross-frequency coupling also demonstrated better external brain tracking of stress rhythm. These results were corroborated by between-subject Pearson’s correlation (English: *r* = 0.170, *p* = 0.017; Chinese: *r* = 0.082, *p* = 0.245). Collectively, these findings highlight the close link between (exogenous) neuro-audio synchronization and (endogenous) neural coupling of delta-theta oscillations coding the stress patterns in speech that are evident exclusively in English speakers who have extensive stress-language exposure.

**Figure 7:**
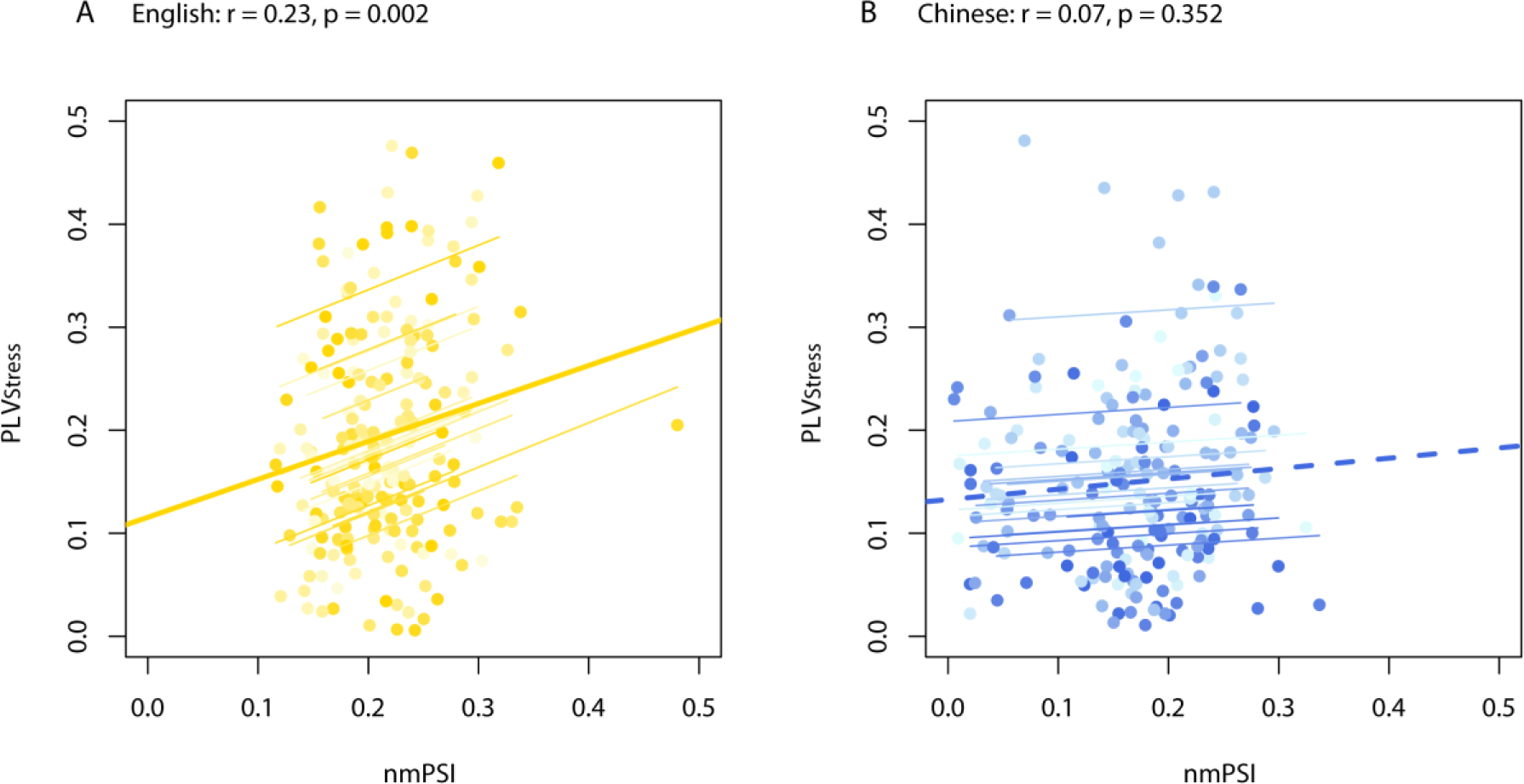
Repeated measure correlations between internal cross-frequency coupling and external audio-neuro tracking in (A) English speakers, and (B) native Mandarin-Chinese speakers. PLV_Stress_ refers to neural-to-stress rhythm phase locking; nmPSI represents the phase coupling of delta-theta neural oscillations to hierarchical stress rhythms. Dots/thin lines = individual data; solid thick line = significant overall relation; dotted thick line=n.s. relation.

## 4. DISCUSSION

Here, we provided new evidence that neural oscillations across multiple time scales mirror the hierarchical nature of the acoustic stress rhythm in speech and do so in a language-dependent manner. Specifically, analyses of phase synchrony measures revealed five key findings: (i) brain oscillations at multiple temporal scales (delta and theta) concurrently phase locked to the rates of stress and syllable rhythms, (ii) amplitude was a more robust stress indicator than duration; (iii) only English speakers demonstrated enhanced multiscale brain-to-speech tracking at the dominant stress rate (2 Hz) and syllable rate (4 Hz) characteristic of natural English, while this phenomenon was absent in Chinese speakers; (iv) both English and Chinese individuals showed delta-theta phase coupling within the brain that mirrors the stress-syllable hierarchy in natural speech but such coupling was stronger in native English listeners; (v) individuals with superior nesting of neural oscillations (i.e., English listeners) also showed enhanced cortical-acoustic tracking to stress. Collectively, our findings suggest brain entrainment mechanisms coding aspects of speech-language are not solely acoustic-induced responses but benefit from phonological knowledge gained from sustained experiences of speaking and listening to a stress-dominant language.

### 4.1 Cortical encoding of stress rhythm via delta phase-locking depends on language experience

Prominent oscillatory-based models (e.g., TEMPO, AST) of language temporal processing have generally overlooked the delta band of the EEG which corresponds to slower-than-syllable rhythms (Ghitza, 2011; Ghitza & Greenberg, 2009; Hickok & Poeppel, 2007; Poeppel, 2003). Our PLV_Stress_ findings show delta oscillations phase-lock to slower (< 4 Hz) acoustic regularities, explicitly tagging the stress rhythms in English. Notably, such neural-audio synchronization is modulated by various acoustic attributes (i.e., stress rate and cue type) and diminishes in individuals with a foreign language background. Prior studies have assumed delta oscillation retain an analogous role as theta, parsing continuous speech into sequential delta-size chunks (Giraud & Poeppel, 2012; Rimmele et al., 2021). However, critical to our findings is the proposition that delta oscillations are associated with the hierarchical nesting role of stress rhythms. Hence, we propose that delta oscillations serve a higher-order mechanism, extending beyond simple stress segmentation to facilitate temporal integration and establish a cohesive phonological representation. Indeed, this nesting function of delta is confirmed by our cross-frequency phase coupling analysis (i.e., nmPSI), evident during the processing of stress patterns. Presumably, delta oscillations coordinate syllable nesting and stress segmentation to streamline ongoing speech processing. Our premise is particularly compelling given the large numbers of individual syllables in connected speech and the consequent cognitive demands on memory and attention, which are consistent with the increased delta activity in working memory where attention is focused on an internal representation (Bidelman et al., 2021; Harmony, 2013). Moreover, this converges with neuroimaging evidence pointing to lexical and semantic grouping via delta oscillatory activities, even in the absence of acoustic boundary cues (Ding et al., 2016; Lo et al., 2022; Meyer et al., 2017).

Furthermore, we found English listeners exhibited stronger phase encoding of stress patterns compared to Chinese speakers, independent of stress cue, rate, and salience. These findings are in line with previous cross-language, electrophysiological studies which show differential brain responses in native vs. nonnative listeners to stress information (Chung & Bidelman, 2016). English listeners’ superior encoding and tracking of stress patterns could lie in their heightened perceptual sensitivity and detection accuracy of stress patterns (Chrabaszcz et al., 2014; Qin et al., 2017). Conversely, Mandarin speakers’ poorer synchronization to ongoing acoustic cues essential for discerning English stress patterns is likely due to their more limited exposure and motor practice of a stress-dominant language. Such experience-dependent effects emerge in both groups’ EEG. Chinese responses were severely hindered by stress manipulations whereas English responses were largely impervious (Fig. 6). Consequently, Chinese speakers struggle to capture acoustic stress regularities in ongoing speech and subsequent failure to segment delta-size chunks might be due to a “perceptual narrowing” of speech representations that are not behaviorally relevant cues in Mandarin (Jeng et al., 2011; Tierney & Nelson III, 2009). Perceptual narrowing due to synaptic neural pruning could manifest at the macroscopic level in the less synchronized brain-to-speech oscillations we find in our EEG data. This lack of coherence, readily achieved by native English speakers, further suggests delta oscillatory synchronization is not merely a passive “bottom-up” mechanism. Rather, we suggest it is sculpted by “top-down” regulation fostered by a listener’s lifetime of sensory experiences and accumulated phonological stress knowledge inherent to speaking a specific language.

### 4.2 Multilevel brain-to-speech synchronization is optimized for the natural rate of stress rhythms

Research has emphasized the importance of amplitude envelope in the brain’s neural entrainment to speech at the syllable level (Assaneo & Poeppel, 2018; He et al., 2023). Our PLV_syllable_ findings further show that syllabic-theta synchronization is critical to suprasegmental processing of stress. Similar dual-frequency synchronization has been observed in intelligible story listening (Gross et al., 2013; Park et al., 2015) and other cognitive tasks (Palva & Palva, 2018). Such nesting of brain responses might be necessary for stress processing since it simultaneously occurs on two distinct timescales (theta-syllable; delta-stress) and each band could track different frequency-specific acoustic information. However, such architecture does not necessary require there be an exhaustive linear division of the incoming speech signal into individual segments (Ghitza, 2013). Rather, a heterodyning of neural oscillation might help establish hierarchical time resolution windows that synchronize to different features of the input (e.g., syllable vs. stress). Corroborated by our PLV and nmPSI measures, our data converge with prior models of language processing that, at least theoretically, can be described as a series of coupled neural oscillators carrying different features of the linguistic signal (Ding et al., 2016; Ghitza, 2011; Hickok & Poeppel, 2007; Park et al., 2015). Our work extends such frameworks by implicating multi-time resolution and experience-dependent plasticity to these models.

Furthermore, we observed differences in cortical-acoustic synchronization across syllable and stress rates. English (but not Chinese) speakers demonstrated enhanced PLV_Stress_ at 2 Hz, closely aligning with the nominal speed of English stress (Dauer, 1983; Leong, 2012; Tilsen & Arvaniti, 2013). Interestingly, we found a similar phase-locking enhancement at 4 Hz, the dominant syllable rate typical for many languages (Ding et al., 2017; Greenberg et al., 2003; Greenberg et al., 1996; Tilsen & Johnson, 2008), that was evident in English speakers but absent in Chinese speakers. This contradicts previous assertions that cross-linguistic differences in neural-acoustic synchronization only appear at the supra-syllabic (but not syllabic) level (Blanco-Elorrieta et al., 2020; Ding et al., 2016; Rimmele et al., 2023), simply because the latter is similar across languages (Ding et al., 2017). However, our analysis further confirmed that the group differences at syllabic level were exclusively marked at the 4 Hz syllable rate. These findings suggest that neural enhancement at the universal syllable rate (4 Hz) might disappear when processing syllables within a foreign *stress context*. As evidenced by our PLV_Stress_ results, the absence of 4 Hz syllabic enhancements in Chinese speakers presumably results from their limited neural coding of stress patterns at the supra-syllabic level (here 2 Hz). Alignment of brain activity to dominant natural rhythms is the key to observing enhancements in neural-speech entrainment (He et al., 2023). The lack of such effect in nonnative listeners implies that a failure to synchronize with higher-order properties of the speech signal (i.e., stress rhythm) might actually impede essential neural processing at lower levels of the hierarchy (i.e., syllable tracking). Future studies are needed to test this possibility.

### 4.3 Neural coupling of delta-theta oscillations mirrors phase hierarchy between speech rhythms

To empirically test for hierarchical relations between frequency-specific neural oscillations, we measured *n:m* phase synchrony within the EEG, which can be intuitively described as the ongoing phase of *n*-cycles of an oscillation synchronizing with *m*-cycles of another oscillation (Leong, 2012; Schack & Weiss, 2005). Unlike our PLV analysis, which reflects the brain’s tracking of sound features of the external acoustic signal, nmPSI reflects oscillatory coupling internal to the brain (*brain-to-brain* synchronization). Our nmPSI results uncovered significant phase-phase coupling between delta and theta neural oscillations for both English and Chinese speakers^4^, closely mirroring the phase hierarchy carried by acoustic stress and syllable envelopes. Moreover, Chinese listeners demonstrated similar delta-theta coherence as English speakers under high stress salience, which was not observed in external neuro-stress tracking (i.e., PLV). These findings indicate a robust neural hierarchy of delta and theta oscillations, even when listeners are less experienced with stress rhythm. Additional examples of hierarchical coupling stems from studies showing increased delta-theta phase-amplitude coupling during intelligible story perception (Gross et al., 2013). Thus, the existence of such nesting in multiple domains of speech processing suggests delta oscillations might play a higher-order role, reorganizing both the phase and amplitude behaviors of theta oscillators that code different properties of the linguistic signal, stress or otherwise.

Converging with our multilevel PLV results, nmPSI measures also demonstrated hierarchical nesting between neural oscillations. These findings demonstrate that ongoing auditory delta oscillations become synced with the *external* acoustic stress regularities which might then formulate an oscillatory hierarchy *internal to the brain* during speech processing, or vice versa. Supporting this notion, we found significant correlations between nmPSI and PLV_Stress_, indicating that a higher degree of internal hierarchical coherence predicts the external alignment of auditory oscillations with stress patterns, or vice versa. Our findings establish a new, heretofore unrecognized relationship between internal neural coherence and external neural tracking across multiple scales, that also varies in a language-dependent manner.

### 4.4 Amplitude cues dominate the neural encoding of stress

Another aim of our study was to evaluate how different acoustic attributes of stress entrain brain oscillations in native vs. non-native speakers. Though English listeners outperformed Chinese listeners in PLV_Stress_, brain-to-acoustic tracking was generally enhanced for stress patterns carried by amplitude compared to duration cues regardless of group. Additionally, English listeners showed more robust syllable tracking (PLV_syllable_) than Chinese individuals for amplitude cues. These findings imply that amplitude-signaled stress more effectively fosters delta-stress synchronization and, at least in English speakers, improves syllabic neural tracking. In general, our data suggest that amplitude cues are more perceptually salient to distinguish stress patterns for both English and Chinese speakers, consistent with prior studies (Chrabaszcz et al., 2014; Zeng et al., 2022). Furthermore, our findings reinforce the “iambic-trochaic law” in linguistics, which posits an innate tendency for intensity-contrasting elements to be perceived as trochaic stress (Strong-weak patterns)—the characteristic of our stimuli—whereas duration-varying components lean towards iambic perception (Crowhurst, 2020; Hay & Diehl, 2007; Hayes, 1995).

However, cross-frequency coupling within the brain also revealed distinct acoustic preferences between language groups. For English speakers, nmPSI values were invariant to acoustic stress cue type (amplitude ≈ duration) and salience (high ≈ low), indicating remarkable stability in delta-theta brain coherence among native speakers even in scenarios of weak stress cues. Contrastively, Chinese speakers showed significant acoustic-driven effects in nmPSI, with stronger coherence for more salient stimuli and duration vs. amplitude cues. This indicates that their neural coherence induced by (English) stress patterns is perhaps more vulnerable to acoustic variations. However, it is worth noting that for duration-based stimuli, nmPSI in Chinese listeners exceeded the baseline nmPSI to stress-free syllable trains. Speculatively, this implies that even non-native listeners may have attempted to construct an internalized hierarchy for duration-based stress stimuli. A possible explanation may lie in the inherent structure of the Chinese language. Chinese might possess a phonological hierarchy similar to English, but one that is organized by tonal instead of stress rules (Duanmu, 2007; McCawley, 1978). For instance, Chinese syllables that carry lexical tone are longer than their weak neutral (non-tone) counterparts. And a supra-syllabic unit emerges by following the rule that neutral tones only present after those with tones (Duanmu, 2004; Li et al., 2014). Such duration-related phonology in Chinese may transfer as a cue-weighting strategy to process an unfamiliar stress hierarchy (Holt & Lotto, 2006; Zhang & Francis, 2010), leading to effective but diminished delta-theta brain coherence.

### 4.5 Conclusions

Collectively, our data demonstrate an intricate interplay between neural oscillations, speech rhythms, and stress hierarchical phonology, providing a new dimension to our understanding of perceptual speech processing. Our findings bridge several gaps, showing multiple timescales of neural oscillations internally cohere and externally synchronize with syllable and stress rhythm. Crucially, individual variations in hierarchical coherence internal to the brain predict their external entrainment ability, and vice versa, essentially reshaping the brain’s engagement with the rhythmic essence of speech. English speakers displayed native advantages in oscillatory synchrony during stress encoding, emphasizing benefits from “top-down” processing rooted in their lifetime exposure to a stress-dominant language. Our results highlight the critical role of brain oscillations in tracking and encoding stress and syllable rhythms in a language-dependent manner.

## Acknowledgements

Requests for data and materials should be directed to G.M.B. This work was supported by Institute for Intelligent Systems Student Organization Thesis/Dissertation Award funding awarded to D.H. and the National Institute on Deafness and Other Communication Disorders (R01DC016267) awarded to G.M.B.

## Author Contribution

Deling He: Conceptualization; Formal analysis; Investigation; Methodology; Visualization; Writing – original draft; Writing – review & editing.

Gavin M. Bidelman: Conceptualization; Formal analysis; Funding acquisition; Methodology; Supervision; Writing – original draft; Writing – review & editing.

Eugene H. Buder: Conceptualization; Supporting; Writing – review & editing

## Footnotes

1 Leong (2012) coined the term “stress foot”, which is also known as metrical or prosodic foot, to emphasize its holistic role in speech, blending rhythmic segments with suprasegmental features. In the current study, stress foot rhythm, or stress rhythm, is used interchangeably to denote the continuous suprasyllabic rhythm that arises from the process of assigning stress to string together multiple syllables.

2 Due to data logging error, one participant yielded 9 epochs for the condition of amplitude stress cue of low salience at 3 Hz, and another participant yielded in 11 epochs for the condition of amplitude stress cue of high salience at 1 Hz.

3 In the duration condition, while the stimuli preserve the overall syllable rhythm, the duration manipulation between stressed and unstressed syllables inevitably leads to non-isochronous syllables which might create jitter in the response and weaken PLV_syllable_. However, PLV_syllable_ magnitudes were, on average, similar between amplitude and duration-cuing stress (e.g., Fig. 4) suggesting any jitter introduced in our stimuli did not negatively impact PLV_syllable_.

4 The nmPSI responses in Chinese speakers were below baseline under for low salience, amplitude-signaled stress.

