## Supplemental Figure 1 for "Cross-linguistic and acoustic-driven effects on multiscale neural synchrony to stress rhythms"

**Supplemental material**

**Behavioral responses**

**Figure S1** illustrates the correct percentage of stress pattern perception during EEG recording. A mixed model ANOVA showed significant three-way interaction effects of stress cue * salience * group ($F_{1,352}$ = 4.55, $p$ = 0.034, $n_p^2$ = 0.01) and stress cue * rate * group ($F_{2,352}$ = 3.64, $p$ = 0.027, $n_p^2$ = 0.02). Post hoc tests revealed the stress cue * salience * group interaction was attributed to duration-modulated stress rhythm under low stress salience, resulting in higher correct percentages compared with amplitude-signaled stress ($p$ = 0.363) within the Chinese group. Meanwhile, the cue * rate * group interaction resulted from duration-modulated stress rhythm at a rate of 3 Hz, leading to higher accuracy relative to amplitude-signaled stress ($p$ = 0.000) within the English group. However, it is important to recognize that neither interaction was driven by group differences, highlighting that both English and Chinese speakers performed comparably (and near chance level) in behaviorally identifying the stressed syllables (English: 55.93% ± 22.67%; Chinese: 59.30% ± 26.77%). While we hesitate to interpret null findings, a plausible explanation for these floors effects might be that both native and non-native speakers found the task too difficult to perceive the location of stressed syllables (or stimuli were too perceptually ambiguous) or participants did not fully understand the task instructions. Future studies using two-alternative forced choice approaches might also produce more definitive perceptual results.

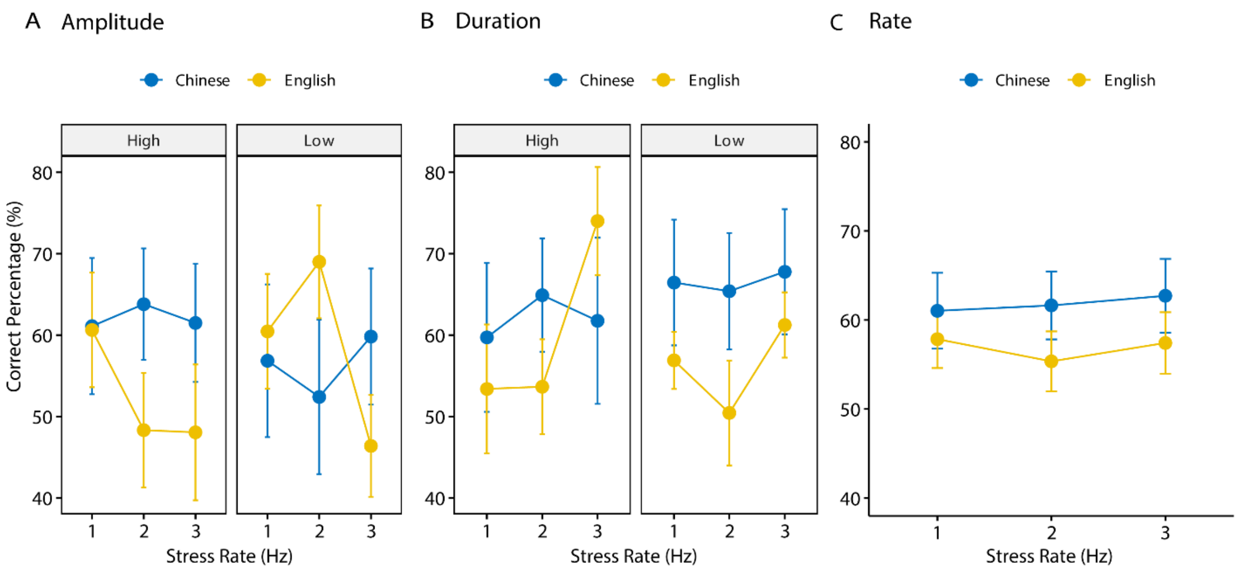


**Figure S1: Group comparisons in correctly perceiving stress patterns by stress rate and salience** modulated by (**A**) amplitude envelope and (**B**) syllable duration. (**C**) no group differences in correct percentage by stress rates with a dataset arrogated across saliences and cues. error bars = ±1 s.e.m.
